# Turn-on Rate Determines the Blinking Propensity of Rhodamine Fluorophores for Super-Resolution Imaging

**DOI:** 10.1101/2022.10.17.512490

**Authors:** Ying Zheng, Zhiwei Ye, Yi Xiao

## Abstract

Live-cell single-molecule localization microscopy has advanced with the development of self-blinking rhodamines. A pK_cycling_ of <6 is recognized as the criterion for self-blinking, yet partial rhodamines matching the standard fail for super-resolution reconstruction. To resolve this controversy, we constructed two typical self-blinking rhodamines (pK_cycling_ = 5.67, 5.35) and a tetramethylsulfonamide rhodamine with unfit pK_cycling_ characteristic (7.00). Kinetic study uncovered slow equilibrium rates and limited blink numbers resulted in the reconstruction failure of partial rhodamines. From the kinetic disparity, a turn-on rate was abstracted to reveal the natural blinking frequency. The new parameter independent from applying laser satisfactorily explained the imaging failure, efficacious for determining the propensity of self-blinking from a kinetic perspective. Following the prediction from this parameter, the tetramethylsulfonamide rhodamine enabled live-cell super-resolution imaging of various organelles through Halo-tag technology. It is convinced that the turn-on rate would be a practical indicator of self-blinking and imaging performance.

## Introduction

Super-resolution imaging technology has revolutionized live cell fluorescence imaging, making it possible to observe the subtle physiological activities of living cells.^[1]^ Among various super-resolution technologies, the single-molecule localization microscope (SMLM) provides one of the highest imaging resolutions.^[2]^ This technique requires molecular switches between dark and bright states, often temporally and spatially isolated during imaging.^[3]^ Thus, high-performance fluorophores are required. Rhodamine derivatives are prominent candidates because they exist in proton-driven equilibria between the leuco spiro (ring-closed) and fluorescent zwitterionic (ring-opened) forms.^[4]^ Such self-blinking rhodamine can achieve sufficient single-molecule signals without the aid of ultraviolet light or redox additives and has become one of the popular strategies for developing SMLM fluorophores.^[5]^

Up to now, the primary criterion for evaluating spontaneous blinking has been pKcycling, that is, pKa values where equal ratios of ring open/closed structures are reached. Urano et al.^[6]^ proposed that for self-blinking rhodamine suitable for SMLM, its pK_cycling_ value should be less than 6. A larger pK_cycling_ value means that a higher proportion of molecules would exist in the ring-opened form at physiological pH 7.4, which might fail to meet the luminescent sparsity required by SMLM. Based on this consideration, they designed a milestone self-blinking Si-rhodamine dye (HMSiR) with a pK_cycling_ value of 5.8. The zwitterionic forms of HMSiR account for a few percent of the total molecules under normal physiological conditions, and sparse single-molecule blinking can be achieved in SMLM imaging without the aid of ultraviolet light or redox additives. Since then, different research teams have successively developed their self-blinking rhodamines following the pK_cycling_ principle. ^[5, 7]^ Liu and Xu et al^[8]^ established a push-pull model to evaluate the feasibility of rhodamines’ spontaneously blinking, in which they indicated that the calculated Gibbs free energy of the zwitterionic state is slightly higher than that in the ring-closed structure, resulting in the sparsity of fluorophores.

Although pK_cycling_ < 6 is a commonly adopted criterion, there is a non-negligible fact that some rhodamine derivatives with their pK_cycling_ values meeting this demand demonstrated insufficient blinks for single-molecule imaging without extra UV activation light (Figure S1, Rh-Gly). This indicates that some factors regulating self-blinking performance have been missed in the criterion of pK_cycling_. As a parameter associated with the structural equilibrium in the thermodynamic steady state, pK_cycling_ helps assess the initial sparsity of the fluorescence-on molecules; however, it cannot indicate the rate of molecular transformation. Therefore, in developing self-blinking rhodamines for SMLM, it is irrational to emphasize pK_cycling_ determination while ignoring the effect of kinetics on the transitions between fluorophore bright and dark states (Figure 1a).

**Figure 1.**
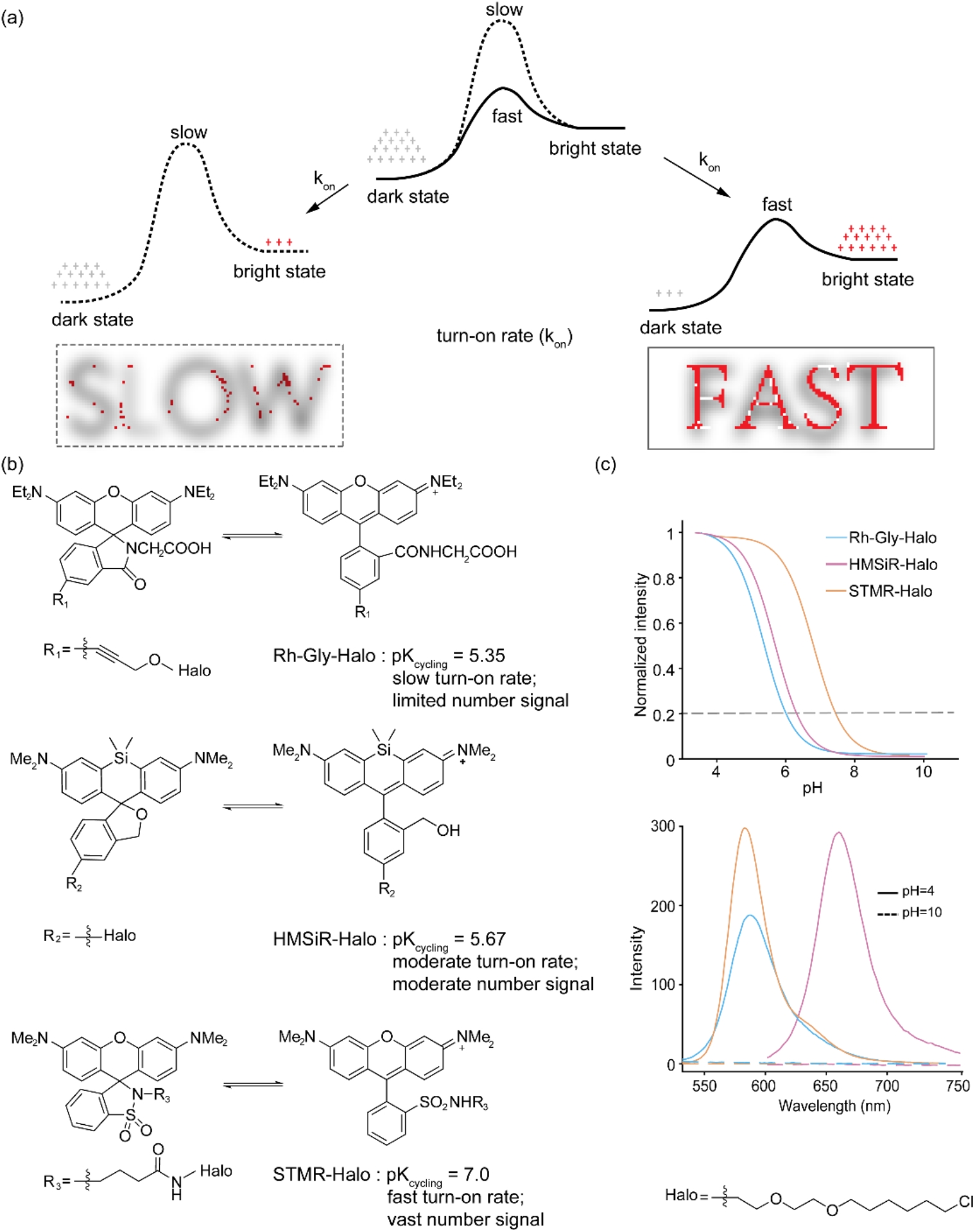
Schematic illustration of equilibrium kinetics on the super-resolution reconstruction. (a) The effect of the turn-on rate of fluorophore in SMLM imaging. (b) Spontaneous spirocyclization equilibrium of rhodamine. (c) Integrated emission intensity vs pH of STMR-Halo, Rh-Gly-Halo and HMSiR-Halo in PBS/C_2_H_5_OH (v/v =7:3) and the spectral comparison in acidic and basic conditions.

This study attempted to mine the properties of self-blinking rhodamine based on kinetic studies and identify the kinetic factors crucial for its super-resolution imaging performance. Therefore, rhodamine dyes with regular and anti-regular pK_cycling_ values for self-blinking were designed and synthesized. Two classic rhodamines, Rh-Gly^[4c]^ and HMSiR, ^[6]^ with self-blinking pK_cycling_ values (5.35 and 5.67, Figure 1c), were used in this study, as the former exhibited irregular super-resolution imaging performance and the latter was the pioneering benchmark self-blinking fluorophore. To further understand the self-blinking mechanism, antiregular tetramethylsulfonamide rhodamine (STMR) was developed with a pK_cycling_ value of 7.00 (Figure S2), which was apparently beyond the preferred region (< 6) for self-blinking rhodamines. STMR has been considered an unqualified self-blinking fluorophore for SMLM, although this has not been confirmed experimentally until now. To facilitate single-molecule and SMLM imaging studies, Halo ligands were installed on all fluorophores to specifically label fused proteins of interest using Halo-tag technology. ^[9]^

Rhodamines were investigated using spectral measurements. It is not possible to directly measure the molecular ring opening or closing rates because the spirocyclization system is in dynamic equilibrium. However, it is possible to record the fluorescence response of ring-opened rhodamines by shifting their equilibrium with pH perturbation. The obtained equilibrium rates (k_E-a_, acid; k_E-b_, base) reflected the degree of molecular ring-opening and ring-closing rates. After acid perturbation, STMR and HMSiR quickly shifted their equilibria at an enlarged fraction of open zwitterions within 40s (Figure 2a and 2e), showing fast acid equilibrium rates (k_E-a_:0.06 s^−1^, STMR; 0.06 s^−1^, HMSiR), whereas the equilibrium of Rh-Gly towards ring-opening zwitterions was slow (> 14400 s, Figure 2c). The acid equilibrium rates of STMR and HMSiR were calculated to be 461 times that of Rh-Gly (k_E-a_ = 1.3×10-4 s^−1^). After basic perturbation, all three dyes rapidly shifted their equilibrium towards the ring-closed leuco form (Figure 2b, 2d, and 2f). The basic equilibrium rates of STMR and Rh-Gly are faster than their corresponding acid equilibrium rates (k_E-b_, 0.33 s^−1^, STMR; 0.17 s^−1^, Rh-Gly). Because k_E-a_ and k_E-b_ indicate the degree of molecular ring-opening and closing speeds, the fast equilibrium rates of STMR and HMSiR suggest that these fluorophores rapidly switched between their zwitterionic and ring-closed forms. In contrast, Rh-Gly with a slow acid equilibrium rate makes it difficult to supply zwitterionic bright states through dynamic equilibrium in super-resolution imaging. It is inferred that this weakness leads to insufficient localization for super-resolution imaging.

**Figure 2.**
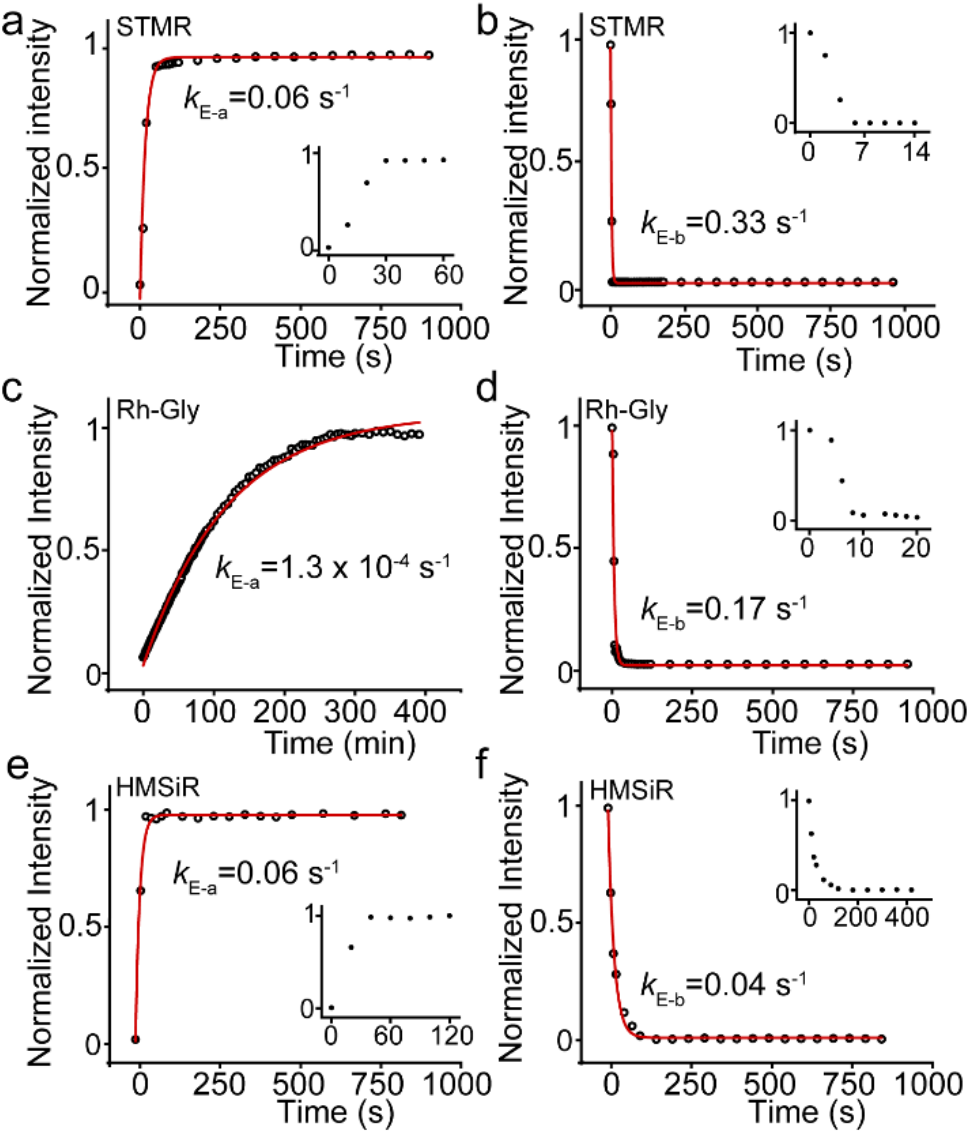
Ensemble kinetic study of spirocyclization equilibrium after acid or basic perturbation. Peak emission intensities (582 nm, 589 nm and 668 nm) are plotted as a function of time, showing the rates reflecting equilibrium shifts to ring-opening (k_E-a_, a, c and e) and ring-closing states (k_E-b_, b, d, f).

To further investigate self-blinking kinetics, the single-molecule characteristics of STMR, Rh-Gly, and HMSiR were studied. The fluorophores were bound to the Halo protein to accurately reflect their kinetics on the tagged protein surface. The obtained fluorophore protein conjugates were further diluted to sparsely adsorb onto the surface of the coverslips, avoiding the overlapping of signals (Figure 3a). Although the three fluorophores were diluted to the same concentration, Rh-Gly exhibited negligible single-molecule signals across the imaging time (Figure S3 and S4). This was probably due to the slow molecular ring-opening transformation of Rh-Gly. Therefore, we increased the concentration of Rh-Gly by 500 times to collect sufficient singlemolecule signals. The single-molecule characteristics of the three fluorophores were evaluated in PBS at pH 7.4, which mimics the physiological environment. The typical fluorescence trajectories obtained for the three fluorophores are shown in Figure 3b and 3c (at 500 W/cm^2^, unless stated otherwise). Although Rh-Gly and HMSiR is “self-blinking” according to their pK_cycling_ values, their molecular fluorescent patterns are dissimilar. Rh-Gly exhibited one dark-to-bright transform, whereas HMSiR exhibited many blinks. Another interesting phenomenon is the large number of blinks for STMR among the three fluorophores, even though this fluorophore shows a pK_cycling_ non-suitable for self-blinking. This discrepancy is consistent in the blink number statistics (54.8 ± 5.1, STMR vs 10.6 ± 1.2, HMSiR vs 2.55 ± 0.03, Rh-Gly), and the order of blink numbers remains under a series of excitation laser powers (Figure 3c and 3d). Other significant single-molecule characteristics are summarized in Figure 3c. All the fluorophores exhibited sufficient single-molecule brightness (152 ± 9, STMR vs. 220 ± 24, HMSiR vs. 96 ± 3, Rh-Gly) for localization. The dwell times of the on-state (on-time) for the three fluorophores demonstrate an order (Rh-Gly, 651 ± 30 > HMSiR, 122 ±10 > STMR, 28 ± 1.2; units are ms), which is consistent with their ensemble equilibrium rates. This consistency suggests that the slow ring-opening/closing transform kinetics might contribute to the dwelling of on zwitterionic states, yet how this ground state equilibrium is involved in the on state with cycles between the ground and excitation states is unknown.

**Figure 3.**
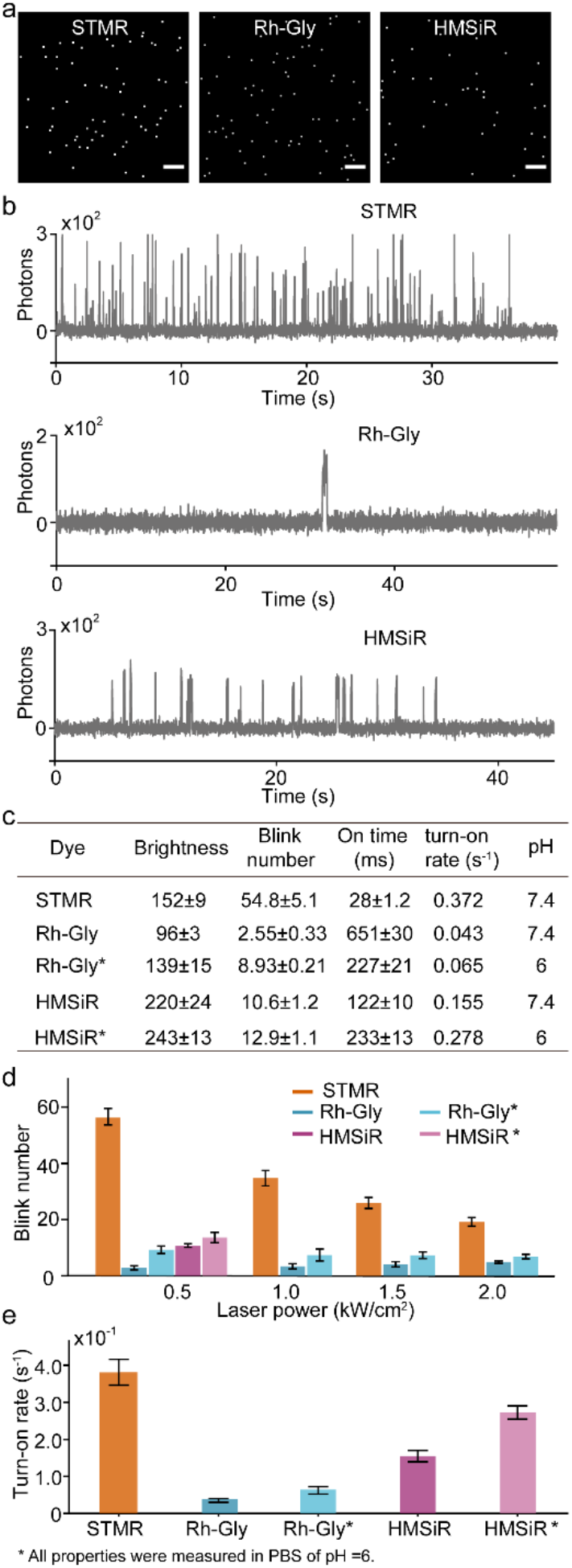
Single-molecule photophysical studies of STMR, Rh-Gly and HMSiR bounded to Halo proteins. (a) Max projection view of imaging stacks shows the sparse distributed single-molecule fluorescent signals. Typical fluorescence trajectories (b) and the summary of single-molecule characteristics (c) of three rhodamines under continuous irradiation of a 500 W/cm^2^ laser. (d) Blink number of three dyes under various laser power. (e) The turn-on rate of STMR, Rh-Gly and HMSiR under 500 W/cm^2^ irradiation. Scale bars: 5 μm.

The single-molecule study above indicated that a limited blink number is the core contributor to the imaging deficiency of Rh-Gly. However, it was not possible to deploy this parameter to identify self-blinking. A significant drawback of this blink number is its dependence on the excitation power, which makes it unrepresentative. Therefore, the molecular transformation kinetics were investigated to uncover an inherent kinetic parameter representing self-blinking. It should be noted that during the single-molecule experiment, a large number of molecules were initially in the dark state and gradually switched to bright states. These dark-state rhodamines are probably in their ring-closed leuco forms in the ground state, and thus, the first transition to the bright state is likely to be the structural ring-opening transformation. Hence, the rate of this first conversion, that is, the turn-on rate, could uniquely represent the molecular ring-opening transform speed, revealing self-blinking kinetics (Figure 3e). Both STMR and HMSiR exhibited rapid turn-on rates (k_turn-on_ = 0.372 s^−1^, STMR; 0.155 s^−1^, HMSiR), showing high probabilities for dark-to-bright transforms in a few seconds. In contrast, Rh-Gly shows a slow turn-on rate, k_turn-on_=0.043 s^−1^, which results in a low probability of observing bright states during acquisition. The orders of turn-on rates (STMR>HMSiR>>Rh-Gly) are consistent with those of blink numbers and equilibrium rates for the three fluorophores, confirming the representativeness of this new parameter for blinking kinetics. The rapid turn-on rates of STMR and HMSiR indicate that these fluorophores only require a low concentration to obtain a sufficient number of single-molecule signals, whereas the concentration of Rh-Gly needs to be significantly increased to enlarge the base number of molecules to obtain sufficient single-molecule signals. Thus, imaging with Rh-Gly should be performed with a high labeling density, which is difficult to obtain and questionable for interference with metabolism in living-cell studies. More importantly, analysis of the turn-on rates under different laser intensities showed that this parameter did not change with the intensity of the applied laser (Figure S5). This independence indicates that this turn-on rate is not influenced by the excitation and can therefore be regarded as a ground-state kinetic parameter. Overall, the turn-on rate is consistent with the blink kinetics and is independent of the excitation states, making it a kinetic parameter for defining the self-blinking level of rhodamines. Although STMR has irregular pK_cycling_ for self-blinking, its high turn-on rate suggests that this fluorophore can spontaneously generate blinks rapidly, predicting a high potential for living-cell super-resolution imaging.

Because the above studies were performed at the same physiological pH, there is a concern that the rate discrepancy originates from the unmatched proton sensitivities of rhodamines, as indicated by their different pK_cycling_ values. It is interesting that structurally different rhodamines exhibit similar turn-on rate orders when their proton sensitivities are equalized by adjusting the environment pH. Hence, the single-molecule properties of Rh-Gly and HMSiR were further investigated at pH 6, at which those fluorophores demonstrate brightening ratios of 0.2 matching that of STMR under physiological conditions (Figure 1c). Although the blink numbers for HMSiR (12.9 ± 1.1) and Rh-Gly (8.93 ± 0.21) are enlarged with the increased proton concentration, these data do not exceed the blinking number of STMR at pH 7.4 (Figure 3c and 3d). Consistently, the turn-on rates of Rh-Gly (0.065 s^−1^) and HMSiR (0.278 s^−1^) also increased but were below the rate of STMR (Figure 3e). Through environmental pH adjustment according to the pK_cycling_ values, the three fluorophores exhibited the same brightening ratios and showed similar proton sensitivities. However, under these conditions, the three fluorophores demonstrated unequal blink rates, which suggests that the pK_cycling_ values could not predict the blink kinetics. In addition, the order of turn-on rates for the three fluorophores is consistent with those acquired under physiological conditions, indicating that this parameter reflects the kinetics inherited from the fluorophore structures.

The pK_cycling_ of HMSiR and Rh-Gly could possibly guarantee static signal sparsity in the context of an appropriate molecular density; however, the ring-opening/closing rates impacted the blinking kinetics defining the self-blinking behavior. HMSiR exhibits a rapid turn-on rate for self-blinking, whereas the slow turn-on rate of Rh-Gly renders it unable to supply sufficient blinking signals for reconstruction. Although the thermal steady pK_cycling_ of STMR does not provide the initial signal sparsity for superresolution imaging, from the perspective of kinetics, a rapid turn-on rate and short on-time of STMR would assist in the rapid formation of a large number of blinking events during imaging (Figure 3). In summary, pK_cycling_ is not suitable for reflecting blinking kinetics, and the ring-opening/closing rates would determine the self-blinking behavior. Thus, the directly measurable turn-on rate can be deployed as a self-blinking indicator that binds the super-resolution imaging performance of self-blinking rhodamines.

To further prove the self-blinking predictions of STMR, we deployed this fluorophore in living-cell super-resolution imaging studies. Through Halo-tag fusion protein technology, STMR was successfully labeled with various structural proteins to mark organelles, including H2B (nuclear histones), Sec61β (endoplasmic reticulum), βTubulin (microtubules), and TOMM20 (mitochondrial outer membrane). Although the microenvironments of various proteins are dissimilar with different hydrophilicities ^[10]^ and pH (e.g., pH ≈ 8.0 for mitochondria, and pH ≈ 7.2 for endoplasmic reticulum, cytosol, and nucleus),^[11]^ during imaging, STMR exhibits similar blinking behavior with a rapid turn-on rate and a large blinking number matching its single-molecule results. Beneficial from the boost blinking kinetics of STMR, living-cell superresolution imaging of nuclear histones, endoplasmic reticulum, microtubules, and mitochondrial outer membrane was successfully achieved without any other additions. The reconstruction imaging exhibits a significant enhancement of resolution compared to the wide-field images (Figure 4 insets and Figure S6-S8). Imaging revealed the nanodomain of H2B proteins in the nucleus, the rod and sheet behaviors of the endoplasmic reticulum (the sheet morphology might be a result of the moderate temporal resolution of single-molecule localization super-resolution imaging),^[12]^ the bending and twisting of microtubules, and the outer membrane boundary of worm-shaped mitochondria.

**Figure 4.**
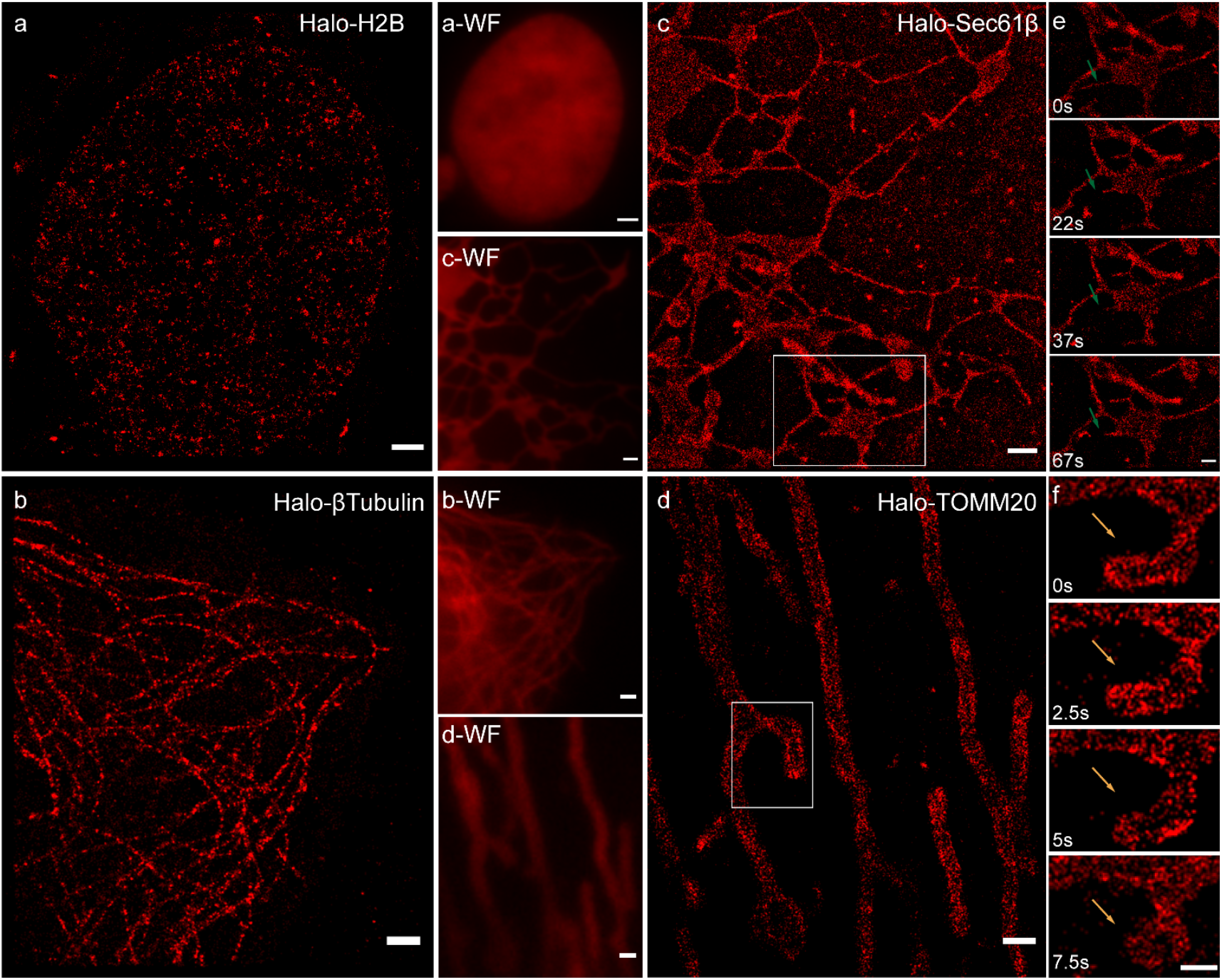
Super-resolution imaging of living cells labelled with STMR through Halo tag technology. Super-resolution reconstruction reveals the distribution of nucleus H2B (a), microtubules (βTubulin, b), the morphology of endoplasmic reticulum (Sec61β, c) and mitochondrial outer membrane (TOMM20, d) proteins in live HeLa or Vero cells. The insets (overlaid with labels suffixed with WF) represent the wide field images from the same location for the corresponding reconstruction. Time lapse images of endoplasmic reticulum (e) and outer mitochondrial membrane dynamics (f). Scale bars: 1 μm (a-d); 0.5 μm (e, f).

In addition, the rapid blinking kinetics of STMR provide many single-molecule signals and a high temporal resolution for reconstruction. This boosting performance enables the monitoring of dynamic morphology for endoplasmic reticulum (for > 60 s; 7.5 s resolution) and mitochondria (for > 10 s; 2.5 s resolution) in living cells. The rod-stretching behavior of the endoplasmic reticulum is shown in Figure 4e, as a small tube (marked by green arrows) gradually shrinks in 22 s and then swings with slight lengthening in the next 40 s. The dynamic morphological evolution of mitochondria is presented in Figure 4f, which shows a shrunken and expanded node. Using STMR-Halo, high-quality single-molecule super-resolution imaging and tracking of different organelles have been achieved in living cells.

In conclusion, pK_cycling_ < 6, as a traditional definition for self-blinking, is misleading, as Rh-Gly with a suitable pK_cycling_ (5.35) fails super-resolution reconstruction. To investigate the inherent kinetics of self-blinking, three rhodamines (Rh-Gly, HMSiR, and STMR) with spirocyclization equilibria were prepared and their ensemble spectral and single-molecule characteristics were studied. The deficiency of Rh-Gly for super-resolution imaging was successfully explained by its kinetics in comparison to HMSiR with a similar pK_cycling_ (5.67); the former rhodamine exhibited a low acid equilibrium rate and blink number, whereas HMSiR showed the opposite. Meanwhile, STMR, with 7.00 pK_cycling_, beyond the traditional self-blinking definition, showed the highest equilibrium rate and blinking number among the three fluorophores, even when the pH was adjusted to equalize the proton sensitivities of the other two fluorophores. Therefore, pK_cycling_ did not indicate self-blinking as it merely reflected the equilibrium balance in the thermal steady state and failed to reveal the blinking kinetics.

To define the self-blinking behavior of rhodamines, a turn-on rate was proposed by mining single-molecule kinetics. STMR demonstrated the most rapid turn-on rate, 9-folds than that of Rh-Gly, 2-folds than that of the self-blinking benchmark HMSiR, consistent with the comparison of the results of ensemble equilibrium rates and single-molecule blink numbers. Most importantly, distinct from the blink numbers, the turn-on rate was unaffected by the excitation laser, proving its relevance to ground-state kinetics and making this parameter a solid indicator for self-blinking. Following the imaging functionality prediction from this parameter, STMR exhibited qualified blinking behaviors to reconstruct the distribution of nuclear H2B proteins, the morphology of microtubules, and transform dynamics of the endoplasmic reticulum and mitochondrial outer membranes in living-cell super-resolution imaging. The identification of the turn-on rate provides an indicator for defining self-blinking rhodamines and bounding super-resolution imaging performance. It is expected that this new kinetic parameter will reset the standard for developing self-blinking rhodamines for future single-molecule super-resolution.

## Acknowledgements

This work was supported by the National Natural Science Foundation of China (Nos. 22174009 and 22004011), the China Postdoctoral Science Foundation (No. BX20200073 and 2020M670754), the Fundamental Research Funds for the Central Universities (Nos. DUT22LAB608, DUT20JC39 and DUT21YG126), and the Dalian Science and Technology Innovation Fund (No. 2020JJ25CY014). Fluorescent imaging was performed with the support of the Chemical Analysis and Research Center at Dalian University of Technology.

---

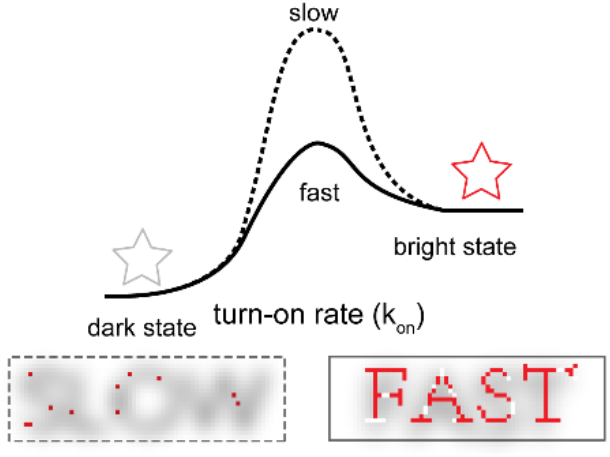

Self-blinking rhodamines meet the demands of single-molecule localization super-resolution for sparse blinking of fluorophores. Turn-on rate is a blink kinetics parameter that predicts the ability of the fluorophore to blink during imaging. The fast turn-on rate can obtain sufficient fluorescence signals during the imaging time to achieve sufficient reconstruction, whereas the slow turn-on rate can only obtain very few fluorescence signals in a short time and allows for the reconstruction of details.

## Notes

### Competing Interest Statement

The authors have declared no competing interest.

## Reference

[1] a) S. J. Sahl, S. W. Hell, S. Jakobs, Nat. Rev. Mol. Cell Biol. 2017, 18, 685; b) Y. M. Sigal, R. Zhou, X. Zhuang, Science 2018, 361, 880.

[2] E. Betzig, G. H. Patterson, R. Sougrat, O. W. Lindwasser, S. Olenych, J. S. Bonifacino, M. W. Davidson, J. Lippincott-Schwartz, H. F. Hess, Science 2006, 313, 1642.

[3] a) T. Ha, P. Tinnefeld, Annu. Rev. Phys. Chem. 2012, 63, 595; b) J. Vogelsang, C. Steinhauer, C. Forthmann, I. H. Stein, B. Person-Skegro, T. Cordes, P. Tinnefeld, Chem. Phys. Chem. 2010, 11, 2475.

[4] a) L. D. Lavis, Biochemistry 2017, 56, 5165; b) A. N. Butkevich, M. L. Bossi, G. Lukinavičius, S. W. Hell, J. Am. Chem. Soc. 2019, 141, 981; c) Z. Ye, H. Yu, W. Yang, Y. Zheng, N. Li, H. Bian, Z. Wang, Q. Liu, Y. Song, M. Zhang, Y. Xiao, J. Am. Chem. Soc. 2019, 141, 6527; d) Z. Ye, W. Yang, C. Wang, Y. Zheng, W. Chi, X. Liu, Z. Huang, X. Li, Y. Xiao, J. Am. Chem. Soc. 2019, 141, 14491; e) Y. Zheng, Z. Ye, Z. Liu, W. Yang, X. Zhang, Y. Yang, Y. Xiao, Anal. Chem. 2021, 93, 7833; f) Z. Ye, Y. Zheng, X. Peng, Y. Xiao, Anal. Chem. 2022, 94, 7990; g) L. Wang, M. S. Frei, A. Salim, K. Johnsson, J. Am. Chem. Soc. 2019, 141, 2770; h) G. Jiang, T.-B. Ren, E. D’Este, M. Xiong, B. Xiong, K. Johnsson, X.-B. Zhang, L. Wang, L. Yuan, Nat. Commun. 2022, 13, 2264.

[5)] a) P. Werther, K. Yserentant, F. Braun, N. Kaltwasser, C. Popp, M. Baalmann, D.-P. Herten, R. Wombacher, Angew. Chem. Int. Ed. 2020, 59, 804; Angew.Chem. 2020, 132, 814; b) A. Morozumi, M. Kamiya, S.-n. Uno, K. Umezawa, R. Kojima, T. Yoshihara, S. Tobita, Y. Urano, J. Am. Chem. Soc. 2020, 142, 9625; c) H. Takakura, Y. Zhang, R. S. Erdmann, A. D. Thompson, Y. Lin, B. McNellis, F. Rivera-Molina, S.-n. Uno, M. Kamiya, Y. Urano, J. E. Rothman, J. Bewersdorf, A. Schepartz, D. Toomre, Nat. Biotechnol. 2017, 35, 773.

[6] S.-n. Uno, M. Kamiya, T. Yoshihara, K. Sugawara, K. Okabe, M. C. Tarhan, H. Fujita, T. Funatsu, Y. Okada, S. Tobita, Y. Urano, Nat. Chem. 2014, 6, 681.

[7] W. Chi, Q. Qiao, C. Wang, J. Zheng, W. Zhou, N. Xu, X. Wu, X. Jiang, D. Tan, Z. Xu, X. Liu, Angew. Chem. Int. Ed. 2020, 59, 20215. Angew.Chem. 2020, 132, 20390.

[8] W. Chi, Q. Qi, R. Lee, Z. Xu, X. Liu, J. Phys. Chem. C 2020, 124, 3793.

[9] a) G. V. Los, L. P. Encell, M. G. McDougall, D. D. Hartzell, N. Karassina, C. Zimprich, M. G. Wood, R. Learish, R. F. Ohana, M. Urh, D. Simpson, J. Mendez, K. Zimmerman, P. Otto, G. Vidugiris, J. Zhu, A. Darzins, D. H. Klaubert, R. F. Bulleit, K. V. Wood, ACS Chem. Biol. 2008, 3, 373; b) A. Keppler, S. Gendreizig, T. Gronemeyer, H. Pick, H. Vogel, K. Johnsson, Nat. Biotechnol. 2003, 21, 86.

[10] a) J. Wilhelm, S. Kühn, M. Tarnawski, G. Gotthard, J. Tünnermann, T. Tänzer, J. Karpenko, N. Mertes, L. Xue, U. Uhrig, J. Reinstein, J. Hiblot, K. Johnsson, Biochemistry 2021, 60, 2560; b) H. Haruki, M. R. Gonzalez, K. Johnsson, PLOS ONE. 2012, 7, e37598.

[11] J. R. Casey, S. Grinstein, J. Orlowski, Nat. Rev. Mol. Cell Biol. 2010, 11, 50.

[12] A. G. York, P. Chandris, D. D. Nogare, J. Head, P. Wawrzusin, R. S. Fischer, A. Chitnis, H. Shroff, Nat. Methods 2013, 10, 1122.

