## Supplemental for "Turn-on Rate Determines the Blinking Propensity of Rhodamine Fluorophores for Super-Resolution Imaging"

Supporting Information
©Wiley-VCH 2021
69451 Weinheim, Germany

Turn-on Rate Determines the Blinking Propensity of Rhodamine Fluorophores for Super-Resolution Imaging

Ying Zheng^†^, Zhiwei Ye^†^,*, and Yi Xiao*

DOI: 10.1002/anie.2021XXXXX

1. Synthetic method

1.1 General

All regents, e.g. MeOH, DMF, acetonitrile, triethylamine, etc., were purchased from commercial suppliers and used as received. Column chromatography was performed with silica gel (200-300 mesh and 300-400 mesh).

1H NMR and 13C NMR were measured on Bruker Avance II 400, Bruker Avance III 500 and Vari-an MERCURY 400 spectrometers. Mass spectra and high-resolution mass spectra were recorded on HP 1100 LC-MSD, gas chromatography/TOF Mass, thermo Scientific LTQ Orbitrap XL, Waters Syn-apt G2-Si HDMS and UPLC/Q-TOF Mass spectrometers.

1.2 Synthesis of STMR-COOH

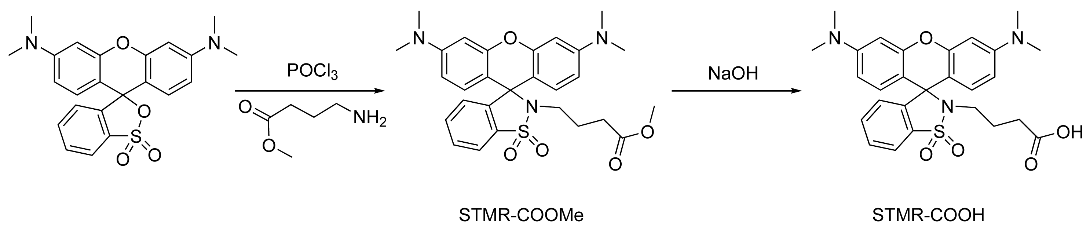

STMR-COOH: Tetramethyl sulforhodamine (STMR) (100 mg, 0.237 mmol) was dissolved in dry 1,2-dichloroethane (5 mL) and stirred vigorously. Phosphorus oxychloride (25 μL, 0.26 mmol) was added at room temperature for 5 min. Then the solution was refluxed for 2h. The obtained mixture was cooled for the next step. The mixture of crude acid chloride was added to a DCM/MeCN (5mL/5 mL) mixture solution of 4-amino-butyricacimethylester (40 mg, 0.26 mmol) and triethylamine at ice bath. After stirring 2h, the crude product was purified through silica gelcolumn chromatography wth a mixture of petroleum ether and ethylene acetate (5:1), v/v) as eluent. STMR-COOMe was obtained as a colorless powder (85 mg, yield 69%). STMR-COOMe (85mg, 0.16 mmol) and NaOH (33mg, 0.81 mmol) was dissolved in methane/H2O (8mL/2mL) and refluxed 3h. After completion of the reaction (monitored via thin-layer chromatography), the solution was evaporated in vacuo. STMR-COOH was obtained through silica gel column chromatography with a mixture of dichloromethane and methane (10:1, v/v) as eluent. The collected solution was dried in vacuo to afford STMR-COOH (80 mg, yield 97%). ^1^H NMR (400 MHz, DMSO-d6) δ 8.14 – 7.92 (m, 1H), 7.72 – 7.52 (m, 2H), 6.93 (d, J = 8.7 Hz, 1H), 6.69 (d, J = 8.9 Hz, 2H), 6.52 (dd, J = 8.9, 2.5 Hz, 2H), 6.41 (d, J = 2.5 Hz, 2H), 2.92 (s, 10H), 2.81 (t, J = 7.2 Hz, 2H), 1.99 (t, J = 7.3 Hz, 2H), 1.44 (p, J = 7.3 Hz, 2H). ^13^C NMR (101 MHz, DMSO) δ 174.18, 152.34, 151.71, 145.60, 134.44, 132.97, 130.05, 129.44, 126.55, 120.58, 109.62, 107.49, 98.41, 66.02, 40.60, 31.15, 24.09. HRMS (ESI) *m*/*z* called for C_27_H_30_N_3_O_5_S [M+H]^+^: 508.1901; found: 508.1907 (*z* = 1).

1.3 Synthesis of STMR

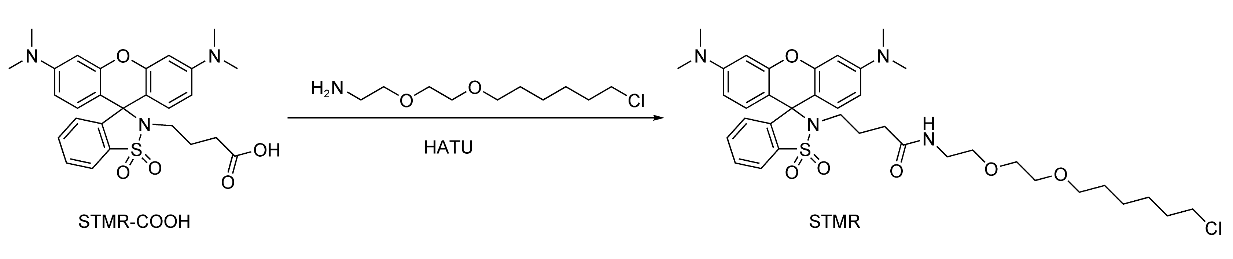

STMR: STMR-COOH (50mg, 0.098 mmol), Halo-NH_2_ (27mg, 0.12 mmol), and HATU (45mg, 0.12 mmol) were dissolved in dry N,N-dimethylformamide (2 mL) in the presence of N,N-disopropylethylamine (20 μL, 0.12 mmol). The mixture was stirred at room temperature for 1h. The reaction mixture was further washed with brine, dried over Na_2_SO_4_, filtered and evaporated. The crude product was purified through silica gel column chromatography (dichloromethane: methane=30:1). STMR (light pink powder, 45mg, yield 64%) was obtained. ^1^H NMR (400 MHz, DMSO-*d*_6_) δ 8.02 – 7.94 (m, 1H), 7.66 – 7.51 (m, 2H), 6.91 (dd, *J* = 5.9, 2.8 Hz, 1H), 6.73 (d, *J* = 8.9 Hz, 2H), 6.51 (dd, *J* = 8.9, 2.5 Hz, 2H), 6.41 (d, *J* = 2.5 Hz, 2H), 3.61 (t, *J* = 6.6 Hz, 2H), 3.45 (s, 4H), 3.36 (d, *J* = 6.6 Hz, 2H), 3.29 (t, *J* = 6.0 Hz, 2H), 3.07 (q, *J* = 5.8 Hz, 2H), 2.93 (s, 12H), 2.88 – 2.73 (m, 2H), 1.87 (t, *J* = 7.3 Hz, 2H), 1.70 (dt, *J* = 14.3, 6.6 Hz, 2H), 1.48 (p, *J* = 6.5 Hz, 4H), 1.42 – 1.34 (m, 2H), 1.34 – 1.20 (m, 2H). 13C NMR (101 MHz, DMSO) δ 171.62, 152.31, 151.64, 145.71, 134.41, 132.85, 130.00, 129.44, 126.48, 120.58, 109.58, 107.55, 98.40, 70.63, 70.01, 69.85, 69.46, 66.00, 45.83, 40.90, 38.83, 33.03, 32.47, 29.52, 26.57, 25.38, 25.21. HRMS (ESI) *m*/*z* called for C_37_H_49_ClN_4_O_6_SNa [M+Na]^+^: 735.2959; found: 735.2958 (*z* = 1).

2 Super-resolution imaging of Rh-Gly without extra UV activation light

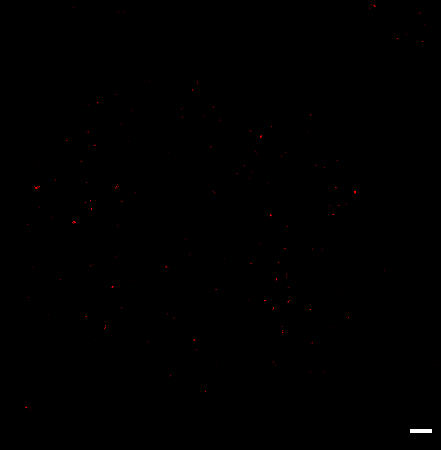

**Figure S1**. Super-resolution imaging of Halo-H2B with 10uM Rh-Gly in a live HeLa cell. Scale bar: 1 μm.

3 Spectroscopic study

Absorption spectra were recorded on Agilent 8453 UV-visible spectrophotometer and fluorescence spectra were recorded on Agilent Cary Eclipse Fluorescence spectrophotometer.

3.1 Method

3.1.1 Calculation of pKa value.

STMR-Halo was prepared as 2 μM solution for measurements. Fluorescence spectra was measured in PBS (100 mM) containing 30% EtOH at various pH values.

3.1.2 Measurement of absorption and emission spectra after binding protein

Fluorophores and Halo protein (excess) were dissolved in PBS (pH=7.4), and the final concentration of the fluorophore was 5 μM. Then the resulting mixture was incubated at 37 ℃ for 2h to obtain a fully reacted solution.

3.2 Spectral Analysis

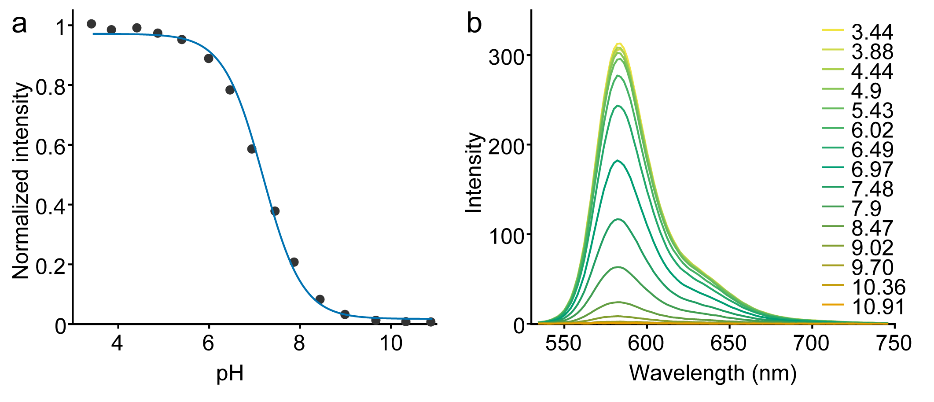

**Figure S****2**. Integrated emission intensity (a) and emission spectral changes (b) vs pH of STMR in PBS/EtOH (v/v = 7: 3).

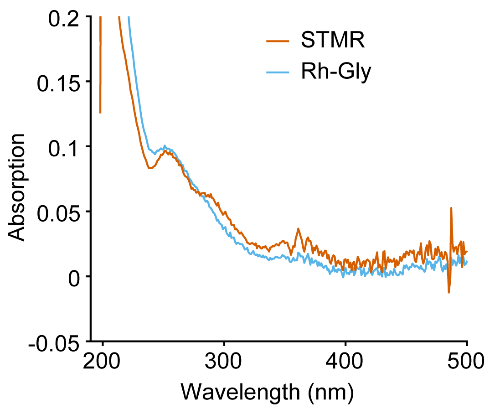

**Figure S3**. Absorption spectrum of the solution diluted 40 times after STMR or Rh-Gly were connected to the Halo-tag protein.

4 Single-Molecule study

4.1 Microscopy

Single-molecule and Super-resolution imaging were studied with a total internal reflection fluorescence microscope (TIRFM) built on an Olympus IX71 inverted microscope as described earlier.^[1]^ The laser light was focused on the back focal plane of a x 100 objective (UAPON 100XOTIRF; 1.49 numerical aperture). An EMCCD camera (iXon DU-897U) was implemented for data acquisition.

4.2 Protein Labeling

STMR or Rh-Gly was labeled to Halo protein in PBS (pH=7.4), then the resulting mixture was incubated at 37 ℃ for 2h. The remaining unbounded fluorophores were removed by protein desalting spin columns (filled with Sephadex G-25 resins).

4.3 Single-Molecule Imaging

The protein solution labeled with STMR (or Rh-Gly) were freshly diluted in PBS at a low concentration to minimize the overlapping between different molecules and transferred to the surface of clean coverslips. The adhesion of proteins to the surface proceed for 30 s and the unbounded proteins were washed out through three times rinses of PBS. The single-molecule fluorescence was recorded with the microscopy described above in conventional mode. The pixel size was 160 nm/pixel and 10000 raw frames were acquired at 10 ms exposure time under different laser power irradiation. At least five measurements were performed for each irradiation condition.

4.4 Single-Molecule Analysis

Raw data were automatically processed with a home-written Matlab software as described in our previous report.^[1]^ Briefly, the molecular candidates were identified by denoising. The spatially ap-proximate molecules were removed during this process. Then the fluorescent trajectories of these molecules were extracted and fitted to a hidden Markov model with a Gaussian distribution as the observation probability distribution.

*Brightness*. Single-molecule brightness in this manuscript was determined as the photon counts from a single molecule during the acquisition time of single frame (10 ms).

*Total collected photons.* In this study, this parameter was calculated as the sum of photons collected from all bright states of a molecule.

*On time.* The time duration of a bright state.

*Blink number.* The counted times of bright-to dark transitions of a molecule.

Signal-to-noise ratio. The signal to noise ratio (SNR) for single intensity trajectory was calculated following the equation below:[XX]

$SNR=\frac{(I_{0}-I_{b})}{\sqrt{I_{0}}}$	(1)

*I_0_* was the maximum pixel intensity of a molecule signal on one frame. *I_b_* was the background noise. The SNR of a molecule was averaged of values from on-states over its entire fluorescence trajectory.

*Duty cycle.* Duty cycle was defined as the ratio between the total time of a single molecule spent on its on-state and the entire imaging time:

$Duty Cycle=\frac{\sum t_{on}}{T}$	(2)

*t*_on_ was the dwell time of a switching event. *T* was the entire imaging time (100 s in this study).

*Turn-on rate.* The rate at which a molecule transitions from the initial dark state to the bright state. (Rhodamine molecules in initial dark state probably exist in their ring-closed form at ground state, as those molecules have not been excited and shown none fluorescence from the beginning.)

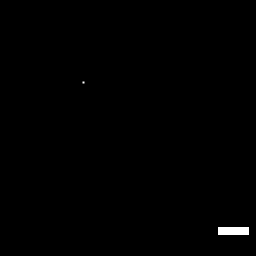

**Figure S4**. Single-molecule fluorescent signals of Rh-Gly at the same dilution as STMR.

**Table S1.** Single-molecule Photophysics of STMR and Rh-Gly

| Dye | Laser Power  (W/cm^2^) | Brightness  (photons/10ms) | Blink number | Duty cycle | On time  (ms) | Turn-on  rate (s^-1^) | pH |
| --- | --- | --- | --- | --- | --- | --- | --- |
| STMR | 500 | 152±9 | 54.8±5.1 | 0.026±0.002 | 28±1.2 | 0.372 | 7.4 |
|  | 1000 | 218±11 | 34.5±5.3 | 0.016±0.002 | 27±0.8 | 0.370 |  |
|  | 1500 | 286±18 | 25.0±2.7 | 0.011±0.001 | 25±1.2 | 0.373 |  |
|  | 2000 | 371±31 | 19.1±5.3 | 0.008±0.001 | 25±0.7 | 0.372 |  |
| Rh-Gly | 500 | 96±3 | 2.55±0.03 | 0.028±0.011 | 651±30 | 0.043 | 7.4 |
|  | 1000 | 147±5 | 2.56±0.36 | 0.014±0.004 | 361±17 | 0.044 |  |
|  | 1500 | 235±9 | 3.15±0.48 | 0.009±0.001 | 186±30 | 0.045 |  |
|  | 2000 | 279±16 | 3.77±0.23 | 0.008±0.001 | 131±19 | 0.045 |  |
| Rh-Gly | 500 | 139±15 | 8.93±0.21 | 0.025±0.011 | 227±21 | 0.065 | 6 |
|  | 1000 | 197±9 | 6.25±1.3 | 0.015±0.004 | 136±45 | 0.067 |  |
|  | 1500 | 282±18 | 6.67±1.0 | 0.010±0.002 | 93±12 | 0.067 |  |
|  | 2000 | 323±10 | 6.18±0.5 | 0.006±0.001 | 63±13 | 0.066 |  |

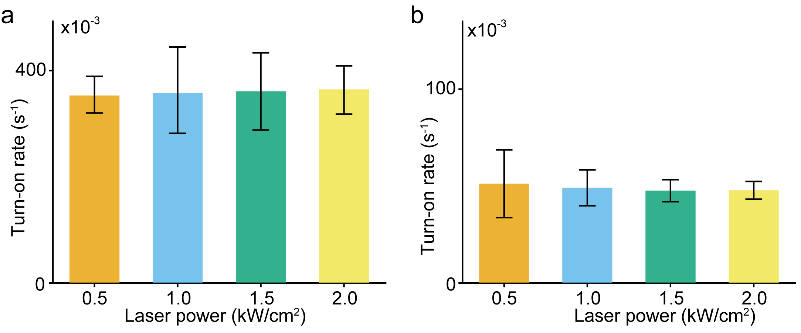

**Figure S5.** The turn-on rate of STMR and Rh-Gly under different laser power in PBS.

5 Super-resolution Imaging

5.1 Cell Culture

HeLa cells were purchased from the Cell Bank of Type Culture of Chinese Academy of Sciences. Vero cells were kindly provided by Procell Life Science&Technology Co.,Ltd. The cells were cultured in full growth medium (cell culture media), that is, minimum Eagle’s medium (MEM) supplemented with 10% fetal bovine serum (FBS, HyClone) and 1% penicillin-streptomycin (PS) solution (x100 HyClone). The culturing condition was humidified atmosphere at 37 ℃ charged with 5% CO_2_.

5.2 Preparation of Cells for Imaging

HeLa cells were transient transfected with Halo-H2B, Halo-Sec61β (Addgene plasmid # 123285), Halo-TOMM20 (Addgene plasmid # 123284), SNAP-ER using Lipofectamine^TM^ 3000 reagent following standard protocol. Vero cells were transient transfected with Halo-βTubulin (Addgene plasmid # 64691) using Lipofectamine^TM^ 3000 reagent following standard protocol. Cells were seeded in cover clips after 24 h transfection.

5.3 Labeling of Fusion Protein

Halo-H2B expressing cells were incubated with 500 nM STMR for 1 h. Halo-Sec61β and Halo-βTubulin expressing cells were incubated with 300 nM STMR for 2 h. Halo-TOMM20 expressing cells were incubated with 100 nM STMR for 2 h. The free remaining dyes were washed with PBS for three times, and were further cultured in CO_2_ incubator with fresh MEM media for 30 min. The imaging media was MEM without phenol red supplemented with 10% FBS.

5.4 Acquisition

A conventional image was acquired with low laser intensity before super-resolution imaging. During super-resolution imaging, a continual 532 nm laser (2 kW/cm^2^) was utilized for excitation. The single-molecule photoswitching signals were recorded at 200 Hz for Halo- Sec61β, Halo-TOMM20, SNAP-ER expressing cells and 25 Hz for Halo-H2B, Halo-βTubulin expressing cells.

5.5 Post-Processing of Super-resolution Imaging Data

Super-resolution imaging analysis was performed in either a ThunderStorm plugin^[2]^ of ImageJ.^[3]^ Briefly, the raw frames were filtered with a difference-of-Gaussians filter to search for signal candidates. Then the point spread functions (PSF) of those candidates were fitted with an integrated form of symmetric 2D Gaussian function (Fitting radius: 3.0 pixel) following Maximum likelihood method^[4]^ to estimate the precise location and single-molecule intensity. The localization precision was calculated according to the Thompson formula.^[5]^ Those PSFs with large widths (> 1.5×median(sigma)), small widths (< 0.5×median(sigma)), and weak intensity (< 100) were eliminated.

Camera readout intensity (I) was converted to photons through the below equations:

$photons= \frac{I\times ADU}{QE\times EMGain}$	(3)

I was the intensity value direct read from camera. ADU was the sensitivity of EMCCD (15.82 electrons per A/D count). EMGain was the gain configuration of the camera (100 in this study). QE was the photon efficiency of the camera.

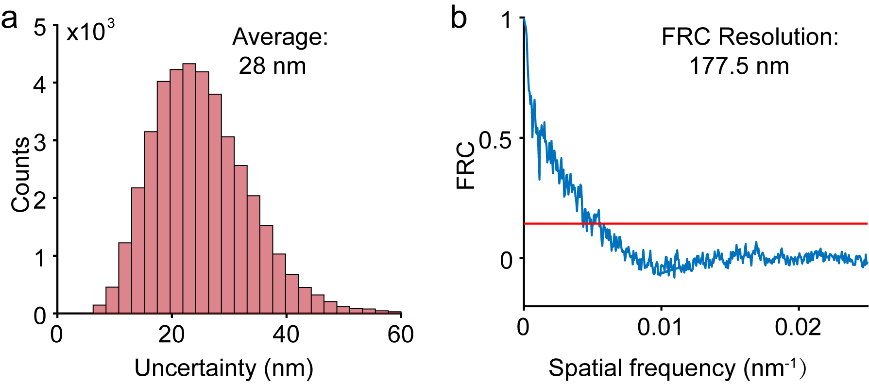

**Figure S6.** Histogram of localization uncertainty (a) and Fourier ring correlation curve analysis of localization (b) in Figure 4a.

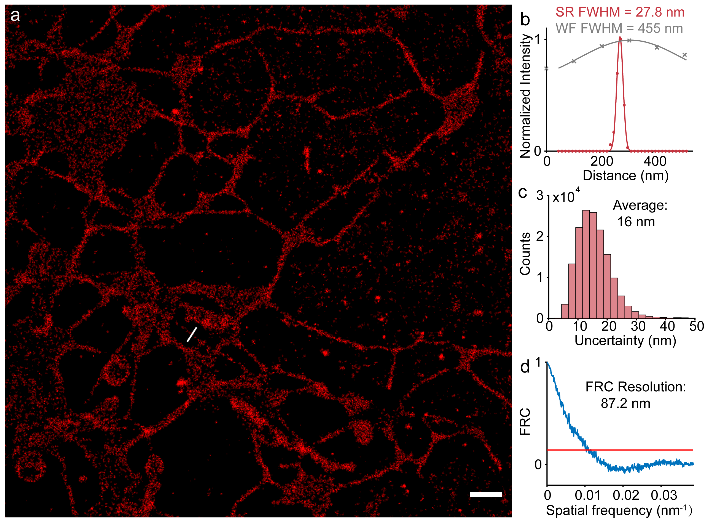

**Figure S7.** (a) Super-resolution imaging of endoplasmic reticulum in a live HeLa cell. (b) Intensity profiles of endoplasmic reticulum tube highlighted with white lines in the super-resolution image (a). Histogram of localization uncertainty (c) and Fourier ring correlation curve analysis of localization (d) of the reconstruction. Scale bar: 1 μm.

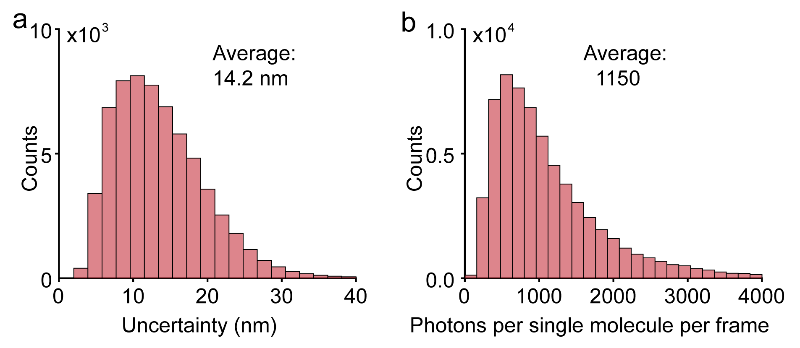

**Figure S8.** Histogram of localization uncertainty (a) and single-molecule brightness (b) in Figure 4d.

6 Characterization spectra

6.1 ^1^H NMR spectrum of STMR-COOH

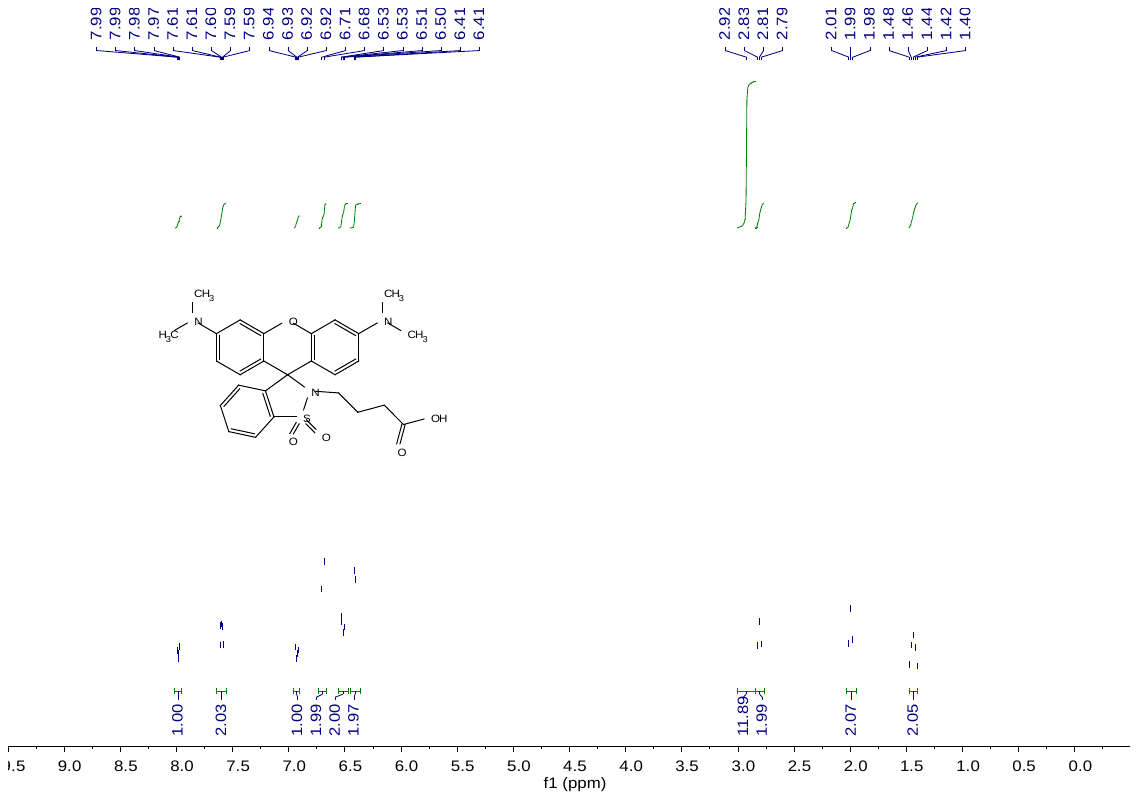

6.2 ^13^C NMR spectrum of STMR-COOH

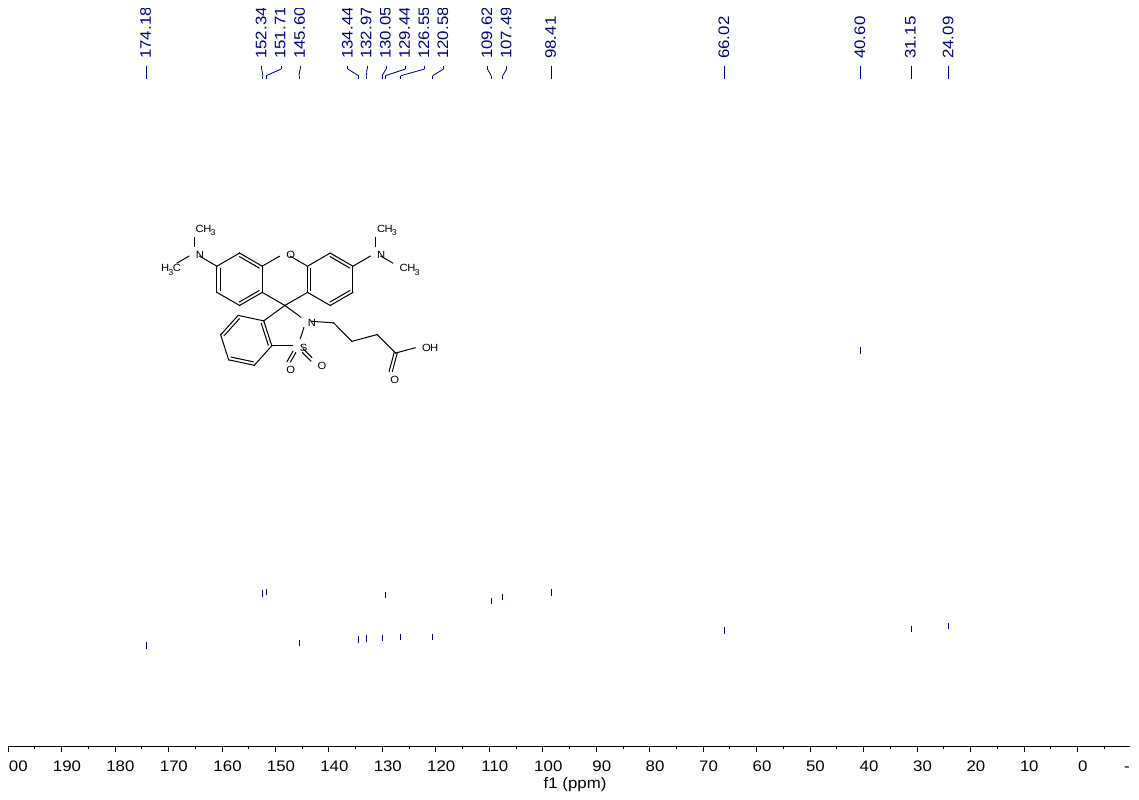

6.3 ^1^H NMR spectrum of STMR

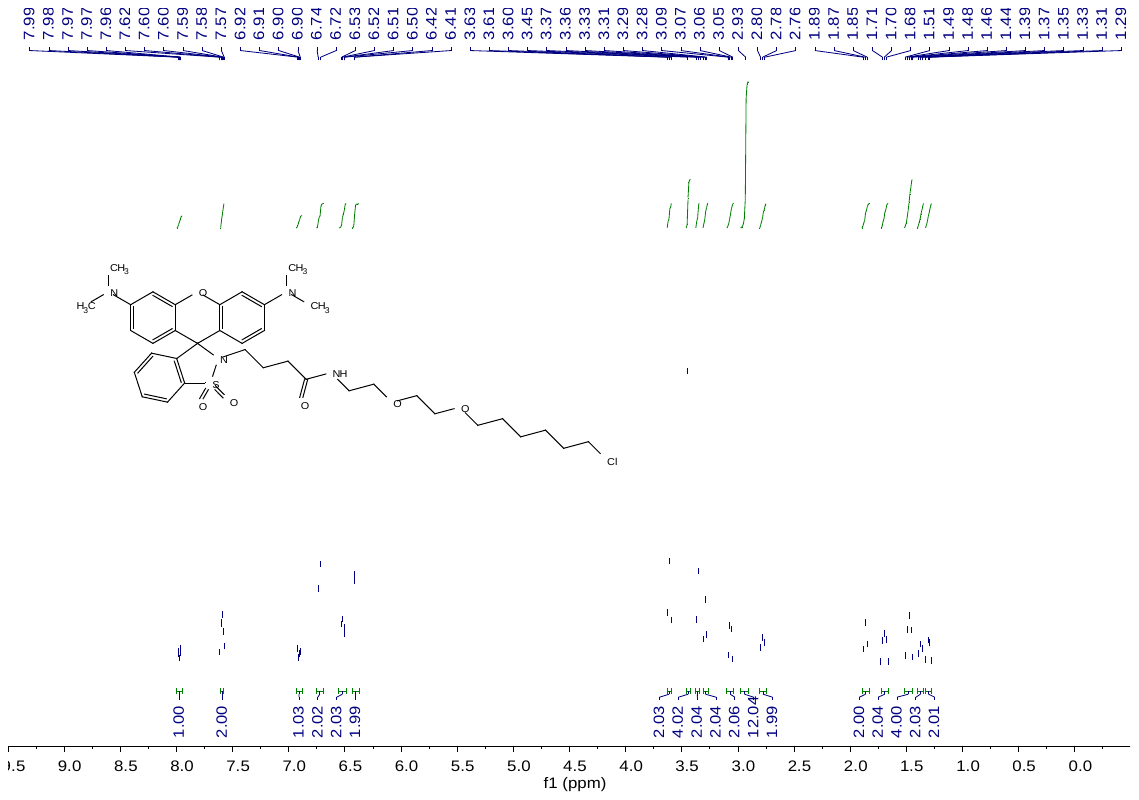

6.4 ^13^C NMR spectrum of STMR

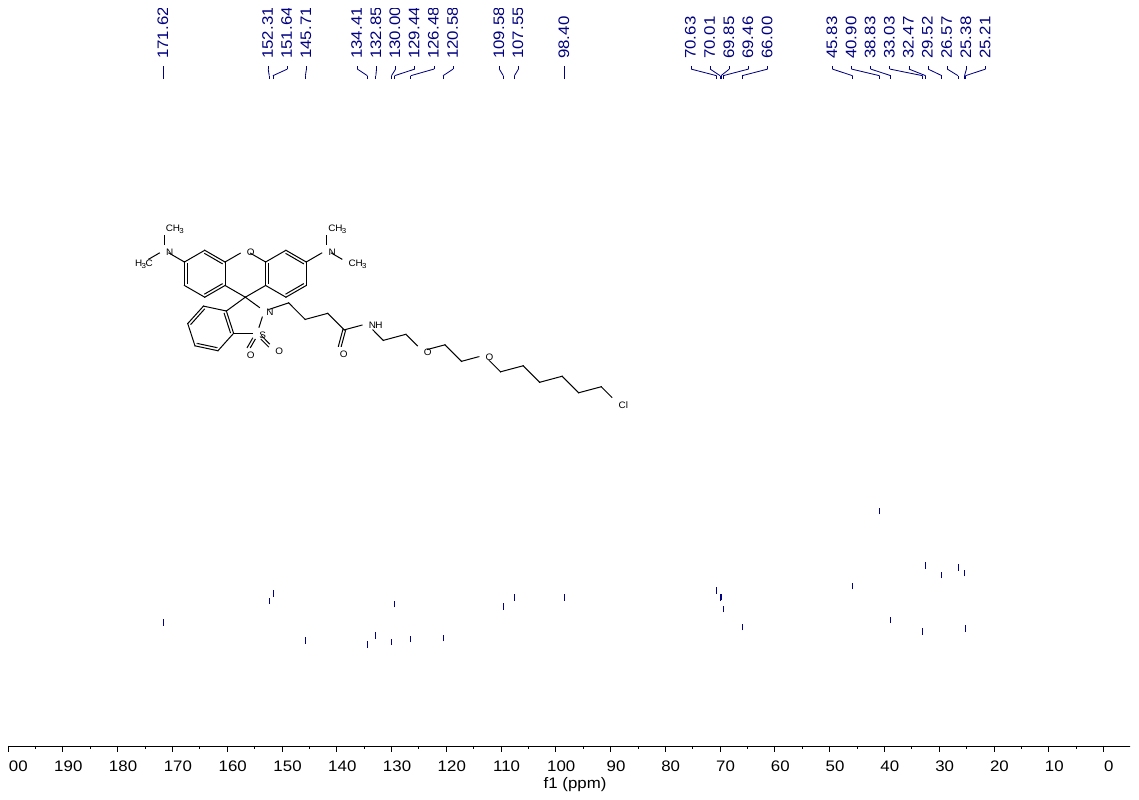
.

### References

[1]	a) Z. Ye, H. Yu, W. Yang, Y. Zheng, N. Li, H. Bian, Z. Wang, Q. Liu, Y. Song, M. Zhang, Y. Xiao, *J. Am. Chem. Soc.* **2019**, *141*, 6527; b) Z. Ye, W. Yang, C. Wang, Y. Zheng, W. Chi, X. Liu, Z. Huang, X. Li, Y. Xiao, *J. Am. Chem. Soc.* **2019**, *141*, 14491.

[2]	M. Ovesný, P. Křížek, J. Borkovec, Z. Švindrych, G. M. Hagen, *Bioinforma.* **2014**, *30*, 2389.

[3]	C. A. Schneider, W. S. Rasband, K. W. Eliceiri, *Nat. Methods* **2012**, *9*, 671.

[4]	a) K. I. Mortensen, L. S. Churchman, J. A. Spudich, H. Flyvbjerg, *Nat. Methods* **2010**, *7*, 377; b) H. Deschout, F. C. Zanacchi, M. Mlodzianoski, A. Diaspro, J. Bewersdorf, S. T. Hess, K. Braeckmans, *Nat. Methods* **2014**, *11*, 253.

[5]	R. E. Thompson, D. R. Larson, W. W. Webb, *Biophys. J.* **2002**, *82*, 2775.

### Author Contributions

Y. Zheng carried out all the experiments, analyzed the data, and wrote the first version of the manuscript. Z.Ye contributed to experiment design, data analysis method, supervised the project and manuscript editing. Y. Xiao supervised the project and contributed to the manuscript writing. All authors have given approval to the final version of the manuscript.
